# digestome: a licence-clean marker-gene panel for functional profiling of anaerobic digestion microbiomes

**DOI:** 10.64898/2026.09.28.755039

**Authors:** Taavi Päll

**Affiliations:** Magrittr OÜ

**Keywords:** anaerobic digestion, biogas, metagenomics, functional annotation, methanogenesis, open data

## Abstract

Functional profiling of anaerobic digestion microbiomes is routinely performed against KEGG or MetaCyc, whose paid commercial licences block fee-for-service analysis for biogas operators, the setting where the results have the most immediate operational value. We present digestome, a curated marker-gene panel for anaerobic digestion built exclusively on sources that are free for commercial use (NCBIfam, public domain; Pfam, CC0; Rhea, CC BY), together with a scorer that reports pathway completeness, branch capability and, for hydrolysis, whether the enzyme is built for export. Benchmarked against 1,401 metagenome-assembled genomes from 134 anaerobic digesters, the panel assigned no methanogenesis route to any of 1,361 non-methanogens, and every acetoclastic call fell within *Methanosarcina* or *Methanothrix*. The result held for 3,043 species representatives from the Genome Taxonomy Database, spanning 203 phyla, which are built differently from binned metagenomes: the only background genomes given a route were two archaea carrying mcrA, and none of the 90 anaerobic methane and alkane oxidisers was given one. Specificity against non-methanogens does not show that a methanogen gets the right route: with a family-level check, 59 of 156 acetoclastic calls in that sample fell on methylotrophic genera that do not use acetate, and a lineage policy removed them, with 14 more left unconfirmed in unnamed genera. Testing whether a catalytic domain shares a polypeptide with an export module reduced the genomes credited with cellulolytic capacity from 597 to 50. A causal graph of the measurement shows that a profile reports the community together with three measurement choices (sampling, DNA extraction and sequencing depth, and reference genomes), which fixes what one profile can attribute and what a prospective validation would have to record.

## Introduction

Industrial anaerobic digestion is commonly regarded as a biological “black box” (De Vrieze, 2020; Koch et al., 2014; Ostos et al., 2024). Biogas plant operators must manage a highly complex microbial community that they cannot directly observe, forcing them to rely on physicochemical proxies, such as volatile fatty acids, the volatile organic acids to total inorganic carbon (VOA/TIC) ratio (also written FOS/TAC or VOA/TAC), ammonium, and biogas composition, that typically register a metabolic disturbance only after a process imbalance has already occurred (Carballa et al., 2015; Koch et al., 2014). Surveys of many full-scale digesters show that community structure reflects process condition (Hassa et al., 2021), yet reviews in the same field still describe microbiome-based monitoring and control as a prospect (Theuerl, Klang, et al., 2019; Theuerl, Herrmann, et al., 2019). Genome-resolved metagenomics is widely proposed as a route to leading, rather than lagging, indicators (Ostos et al., 2024).

Licensing is what first stands between this proposal and a commercial service. The reference resources that streamline functional annotation are tightly restricted for commercial use. For instance, KEGG is not publicly funded; its FTP distribution has been subscription-only since 2011, academic FTP access costs USD 5,000 per year outside Japan, and non-academic use requires a separate commercial licence (Pathway Solutions / Kanehisa Laboratories, n.d.). MetaCyc and BRENDA carry comparable restrictions. Similarly, for carbohydrate-active enzymes, CAZy and dbCAN are non-commercial in their public forms, requiring separate agreements with two independent institutions (AFMB-CNRS & University of Nebraska-Lincoln, n.d.). A consultancy producing a paid report for a biogas plant operator falls squarely within these restrictive terms. Far from a hypothetical constraint on the anaerobic digestion (AD) field, this presents a structural barrier: because published reconstructions of digestion pathways rely heavily on references from KEGG, MetaCyc, and BRENDA, the standard route to a pathway-level analysis is effectively closed to commercial enterprises.

However, the underlying gap is more precise than a sole reliance on KEGG. DRAM pioneered the approach this study follows by condensing individual gene annotations into pathway-level insights at genome resolution (Shaffer et al., 2020). Although DRAM intentionally supports a KEGG-free configuration, its alternative annotation sources for carbohydrate and peptidase function are dbCAN and MEROPS. Because both databases impose their own commercial restrictions, a KEGG-free setup is not yet a commercially clean solution. The true prerequisite is therefore not a KEGG-free pipeline, but one entirely free of licence-encumbered dependencies across the entire stack.

Virtually every tool designed to reconstruct metabolism at the module level from metagenome-assembled genomes (MAGs) relies on KEGG (Table 1), a testament to the maturity of KEGG Modules as a pathway model rather than a shortcoming of the tools themselves. For instance, METABOLIC couples KOfam searches with module context and incorporates motif validation for individual hits. Addressing the inherent challenge of incomplete genomes, MetaPathPredict uses machine learning to predict module presence rather than simply tallying genes. KEMET targets this exact limitation through an approach that closely mirrors our own concerns: it leverages hidden Markov model (HMM) profiles to recover orthologs that standard annotation pipelines miss. Its authors demonstrate that recovering annotations in this manner substantially improves draft genome-scale metabolic models built from MAGs, elevating them in some cases to the accuracy of models derived from complete genomes (Palù et al., 2022). This finding highlights how MAG fragmentation can distort perceived module completeness: the exact failure mode this panel is designed to counteract from the opposite direction. Crucially, KEMET distributes the KEGG licence alongside its source code, so that users inherit its commercial terms transparently.

**Table 1.** Module-level metabolic reconstruction tools and their reference dependencies. Module-level from MAGs: reconstructs pathway modules from metagenome-assembled genomes. Independent of KEGG: runs without KEGG data; where it does not, the KEGG component is named. No restricted dependency anywhere: no reference source in the stack needs a licence for commercial use; restricted sources are named. AD-specific pathway model: pathway definitions written for anaerobic digestion rather than general module definitions.

|  | module-level from MAGs | independent of KEGG | no restricted dependency anywhere | AD-specific pathway model |
| --- | --- | --- | --- | --- |
| DRAM (Shaffer et al., 2020) | ✓ | optional | × (dbCAN, MEROPS) | × |
| METABOLIC (Zhou et al., 2022) | ✓ | × (KOfam) | × | × |
| KEMET (Palù et al., 2022) | ✓ | × (KEGG Modules) | × | × |
| MetaPathPredict (Geller-McGrath et al., 2024) | ✓ | × (KEGG Modules) | × | × |
| this work | ✓ | ✓ | ✓ | ✓ |

The constraint shared by these tools is therefore contextual rather than technical. It becomes restrictive only when applied to commercial work, and it is intrinsically tied to what makes these tools useful: because KEGG Modules serve as the underlying pathway framework, removing KEGG does not merely strip a tool of a database, but leaves it entirely without pathway definitions. Consequently, a licence-clean alternative cannot simply swap out a sequence source; it must supply an independently curated pathway model. Constructing such a model for a single specialised domain, rather than for the entirety of global metabolism, is precisely what makes this problem tractable.

The other obstacle is interpretive, stemming from the fact that marker-gene profiling merely reports what a genome carries, leaving gaps that separate this raw genomic profile (the per-genome calls and abundances for one sample) from the actionable insights a plant operator actually requires. First, a gene may be present, but the pathway it serves may operate in the opposite direction. Second, an enzyme may be encoded but never secreted from the cell; in the context of extracellular hydrolysis, this lack of transport means it never contacts its substrate. Beyond these, a marker for which no detector exists is easily registered as a zero, which is then frequently misconstrued as a complete absence of the pathway. Consequently, translating genomic potential into operational advice remains a fundamental challenge.

Here we address licensing and interpretation together. We describe a curated marker panel for anaerobic digestion drawing only on commercially usable sources, and a scorer built so that each of these gaps produces an explicit statement rather than a potentially misleading number. We then assess specificity against reference genomes of known metabolism and against 1,401 metagenome-assembled genomes from full-scale digesters, and quantify how much the export test changes the hydrolytic capacity attributed to a community.

## Methods

### Panel structure and marker identity

The panel is a tab-separated table of 86 markers across 20 modules (53 core-tier, 33 accessory-tier), covering hydrolysis of carbohydrate, protein and lipid; acidogenesis to acetate, butyrate, propionate, lactate, ethanol, hydrogen and via ethanolamine; acetogenesis including the Wood–Ljungdahl pathway and syntrophic oxidation; and hydrogenotrophic, acetoclastic and methylotrophic methanogenesis, with modules for energy conservation and direct interspecies electron transfer. Every row carries a unique `marker_id` of the form `MODULE:gene`, and all joins between panel, model map and scorer are based on that identifier. Raw gene names are not used as keys. A three- or four-letter gene symbol is unique neither within a panel nor within a reference database, and using one as a join key produces errors that are difficult to see: an EC-based match collapses every model sharing that EC onto a single arbitrary gene name, and a model serving two panel rows can record only one of them, silently scoring the other as absent.

### Marker resolution

Markers resolve to profile HMMs by curated accession, then gene symbol, then EC number. Models are drawn from NCBIfam and Pfam (Mistry et al., 2021). Curated accessions come first: a row can list the accessions of the models that detect it. Marker choice here follows the field where it has been tested directly: formyltetrahydrofolate synthetase outperforms 16S for tracking the acetogenic community in biogas reactors (Singh et al., 2021), and its model is assigned by curated accession in this panel. A row with curated accessions uses those models only: its gene symbol and EC are not indexed at all. Exclusivity is load-bearing rather than tidy: the archaeal A1A0 ATP synthase row matched 120 models through EC 7.1.2.2, an EC that spans every subunit of F-, V- and A-type synthases in both domains. Gene symbol matches are used for rows without curated accessions. This route is not automatically safe: `fmdA` is both the panel’s formylmethanofuran dehydrogenase subunit A and a legitimate symbol for formamidase, an unrelated and common enzyme. EC numbers are the weakest route and are used last. Wildcard ECs (`2.1.1.-, 3.4.-.-`) are excluded entirely; they name an enzyme class rather than a gene, and matching them imported 58 unrelated peptidases, lipases and glycosidases. Of 150 model-to-marker assignments in the current build, 54 are curated, 55 by gene symbol and 41 by EC. Giving curated accessions to a row that a gene symbol already resolved correctly changes no result, as the three methylamine methyltransferases show, but it fixes the assignment so that a later symbol collision cannot silently redirect it. Every non-curated route is auditable: `audit_symbol_matches.py` compares the enzyme the panel requests against the product name of each matched model.

### Score thresholds

Each model is applied at its own curated cutoff (`hmmsearch` from HMMER 3 (Eddy, 2011) with `--cut_nc`) rather than a global threshold, an approach shown to materially affect functional and metabolic inference (Kananen et al., 2025). Across the 144 models, these span 20.8 to 1,400 bits (median 381), so no single threshold is defensible: 94 of the 144 models have a curated cutoff below 450 bits, so a global threshold at that level would reject genuine hits for most of the panel. Pfam and NCBIfam use different conventions. Pfam’s membership threshold is its gathering threshold (GA), which sits above its noise cutoff (NC). For NCBIfam the panel uses NC, the most inclusive of its three curated values (lower than GA for 46 of 127 models). On import each Pfam model’s NC is therefore set to its GA, so one search applies GA to Pfam models and NC to NCBIfam models. The build checks that every pressed model carries a curated cutoff and stops if any does not.

### Pathway gates and taxonomic confirmation

Each pathway gate declares its requirements as groups of alternatives, and gates evaluate to true, false, or not assessable when a required marker has no detector in the database at all. The hydrogenotrophic gate requires `mcrA` and at least four of six C1 pathway steps (`fwdB|fmdB, ftr, mch, mtd|hmd, mer, mtrA`); isoenzymes of one step count once. It also covers methanogens that reduce CO2 with formate. The acetoclastic gate requires `mcrA, cdhA` and acetate activation (`acs`, or `ackA` with `pta`); the methylotrophic gate requires mcrA and a substrate-specific methyltransferase or the shared MT2 step. The acetoclastic and hydrogenotrophic gates, and a methylotrophic call resting on MT2 alone, need a confirming lineage, supplied as a classification against the Genome Taxonomy Database (GTDB) with GTDB-Tk (Chaumeil et al., 2022), because the pathways they read are reversible or shared, and gene content cannot tell the direction. The ACDS/CODH complex serves autotrophic hydrogenotrophs to *fix* carbon rather than to cleave acetyl-CoA, and *Methanothermobacter thermautotrophicus*, an obligate hydrogenotroph, satisfies the acetoclastic gene test. The C1 pathway runs oxidatively in acetoclasts and methylotrophs and in reverse in anaerobic methane oxidisers, all of which satisfy the hydrogenotrophic gene test. The lineage policy is a single table read by the scorer and by report renderers (`assets/acetate_lineages.tsv`), each row with its source. The acetoclastic call is an allow list at genus level: *Methanosarcina, Methanothrix* and *Methanocrinis* (Jetten et al., 1992; Khomyakova et al., 2023; Smith & Ingram-Smith, 2007). For *Methanosarcinaceae*, a family does not suffice, because the family is mostly methylotrophs that do not use acetate (Oren, 2014), so an unnamed genus there is left unconfirmed; that is conservative by design, and it costs sensitivity rather than correcting an error. *Methanotrichaceae* is the one family confirmed as a whole: every described member is an obligate acetoclast, and a genome in an unnamed genus of the family would otherwise have no route at all, since the family is also denied the hydrogenotrophic call below. The hydrogenotrophic call uses a deny list, because hydrogenotrophy is the default for methanogens: the obligately acetoclastic *Methanotrichaceae*, the methylotrophic *Methanosarcinaceae* genera, and the anaerobic methane and ethane oxidisers (Hahn et al., 2020; Haroon et al., 2013; Knittel & Boetius, 2009). A methylotrophic call resting on the MT2 marker alone is accepted only in lineages known to be methylotrophic without a licence-clean substrate-specific marker: *Methanosphaera, Methanomassiliicoccales, Methanomethylicales, Methanofastidiosales* and *Methanonatronarchaeales* (Fricke et al., 2006; Nobu et al., 2016; Sorokin et al., 2017; Vanwonterghem et al., 2016). mcr carriers in anaerobic methane and alkane oxidiser lineages (ANME-1, ANME-2a/b/c/d, *Ethanoperedens, Syntrophoarchaeum*) are counted apart from methanogens. Where taxonomy places a genome outside these rules, the gate is scored absent with the reason stated; where no taxonomy is supplied, or it is too coarse to place the genome, the gate falls back to gene evidence and says so. A genome missing from the taxonomy table is treated as unclassified, never as a negative.

By default, gates are evaluated per genome. As an option, the scorer can require every term of a gate to lie on one contig (contig-level co-occurrence), which keeps the evidence within one organism without needing a bin. We measured what the option costs on the GTDB sample: we scored each methanogen both ways with the final scorer and lineage policy, and took contig counts from the Prodigal gene calls.

### Secretion evidence

The scorer tests whether a catalytic domain and an export module (dockerin, cohesin, carbohy-drate-binding module or S-layer domain) occur on the same polypeptide, which is the architecture of a cellulosome. The test is per-protein; the same two domains in one genome but on different proteins carry no such implication. All five export modules are Pfam families. Signal-peptide prediction was deliberately not used, the standard tool being fee-licensed for commercial work. The three methylamine methyltransferases (mono-, di- and trimethylamine) are assigned by curated accession to pyrrolysine-specific models: PF05369 for the first and the NCBIfam models `dimeth_PyL and trimeth_pyl` for the other two. The corresponding Pfam superfamily spans bacterial homologues that are not methylamine methyltransferases and is not used for detection. All three enzymes carry in-frame amber codons, so all three depend on gene calling that reads through them.

### Reporting non-measurement

A module with nothing measurable reports NA and a status, never `0.0`. Statuses distinguish a marker that is assumed present because it is universal central metabolism, one whose detector is absent from the reference release, one defined only by accessory-tier markers, and one awaiting a licence-clean source. The same rule applies to gates and, through `core_detectable`, to completeness percentages, which must be read against what could be detected rather than against what the panel defines.

### Causal structure of a profile

A profile, as in the Introduction, is the panel’s output for one sample: the route calls and module completeness of each genome, together with each genome’s share of the reads. To state what a profile can and cannot attribute, we drew the causal structure of a digester and of its measurement as a directed acyclic graph (DAG). In a DAG, each node is a variable and each arrow states an assumed direct cause; a missing arrow states that there is no direct effect. The first group of nodes holds the operator’s levers (feed, loading rate, retention time, temperature, trace-element dosing, inoculum, and air dosing for desulphurisation) and the reactor design. The second holds the reactor state (volatile fatty acids, hydrogen, free ammonia, pH) and the trace elements and electron acceptors in the digester. Next come the community and the profile, and three measurement choices that also act on the profile: the sampling point and time, the DNA extraction and sequencing depth, and the reference genomes. The last group holds the other observations and the decision: process data, gas yield, a trace-element analysis, what the operator sees on site, the operator’s next change, and whether a sample is taken at all.

A DAG cannot contain a loop, but the reactor state and the community act on each other. We therefore unrolled the graph into three time slices: the weeks before the sample (about one retention time), the sampling week, and the eight weeks after the profile is delivered. A single edge list holds every arrow with a one-line reason, and the same list draws the figure.

We queried the graph with dagitty 0.3.4 (Textor et al., 2016). We listed the parents of the profile (its direct causes), the ancestors of the community (every node with a directed path to it), and the directed paths from each upstream node to the profile. For the effect of the operator’s next change on the reactor state in the following slice, we listed the back-door paths and the minimal adjustment sets. A back-door path links a decision to its outcome through a common cause rather than through the decision’s effect; an adjustment set is a set of observed variables that blocks every such path, so that the effect can be estimated. Finally, we listed the conditional independencies that the graph implies among observed nodes, that is, pairs of variables that should be unrelated once a stated set of others is held fixed. We tested each one twice, as stated and again conditional on a sample having been taken, because only samples that someone took can be analysed.

The full graph is Figure S1; the script and its output are in manuscript/dag/.

### Software implementation and compute environment

The panel, builder and scorer are command-line tools requiring only HMMER and a Python interpreter, with no third-party Python packages. A Nextflow workflow wraps the analysis path for scale and portability, giving per-genome parallelism, resumability and executor independence, so the same analysis runs on a laptop, a SLURM cluster or a cloud batch service without modification. The workflow invokes the same command-line tools rather than reimplementing them, so neither entry point is privileged: a user with three genomes needs no workflow engine, and a user with fourteen hundred does not hand-roll parallelism. Database staging and panel construction remain scripts, being one-time operations that need outbound network access. All runs used HMMER 3 on 16 cores of an AMD EPYC 7742.

### Validation datasets

#### Reference genomes

Seven reference genomes of known metabolism were chosen to be falsifiable: a metabolic generalist (*Methanosarcina barkeri*), an obligate acetoclastic methanogen (*Methanothrix soehngenii*), two obligate hydrogenotrophs (*Methanoculleus bourgensis, M. thermautotrophicus*), an acetogen (*Clostridium ljungdahlii*), a syntrophic fatty-acid oxidiser (*Syntrophomonas wolfei*), and *Escherichia coli* as a specificity control. *E. coli* carries `pta, ackA, folD, atoB` and the complete *eut* operon, exercising most of the gene-symbol collisions the panel guards against; any methanogenesis gate firing on it indicates a defect. An assertion suite runs the full path (proteome, `hmmsearch`, scorer) over these genomes and checks 24 statements about gates, markers and genus assignment, exiting non-zero on disagreement. One assertion is deliberately inverted: syntrophic butyrate oxidation must report as *not assessable* in *S. wolfei*, an organism that performs the pathway, because two of its four required markers have no licence-clean model. Asserting the honest blank prevents it from silently degrading into a false negative.

#### GTDB representatives

This set tests whether specificity depends on how genomes were constructed. It is 3,043 GTDB r226 species representatives, which are isolate-derived or independently assembled rather than binned from digester metagenomes: every representative of eleven methanogen orders (1,144 genomes, taken whole because they measure sensitivity) together with 1,899 background genomes allocated across 203 phyla in proportion to the square root of phylum size. Proportional allocation would spend the budget on the largest phyla, and equal allocation would weight a singleton phylum like a 36,000-genome one; selection within a phylum is by even stride over sorted accessions, so the sample is reproducible without a seed. The sample was drawn from the GTDB r226 taxonomy and species-cluster files with the sampler as first released; the accession list is distributed with the release, since a later change to the sampler’s order list moves five background genomes. It was classified against GTDB r226. Reports print r232, the release current at the time of writing; every name in the lineage policy was checked to occur verbatim in the r232 taxonomy, and every name the benchmark relies on occurs in the r226 sample, so the policy reads the same under either release. For scoring, one *Methanonatronarchaeales* genome sampled as background counts as a methanogen, and the 90 genomes in anaerobic methane and alkane oxidiser lineages (89 sampled in methanogen orders, one as background) form a class of their own, so the classes hold 1,056 methanogens and 1,897 background genomes (Results). Genes were called with Prodigal. A third of GTDB representatives are assemblies with no NCBI annotation, and the gap falls on uncultured lineages, so downloading proteomes lost 28 phyla outright. Calling genes also keeps this benchmark comparable with the digester catalogue, which was itself Prodigal-called, and it is required for the pyrrolysine marker described below.

#### Digester catalogue

The catalogue comprises 1,401 metagenome-assembled genomes recovered from 134 anaerobic digesters spanning multiple plants, feedstocks and operating temperatures [Campanaro et al. (2020); NCBI BioProject PRJNA602310]. Taxonomic assignments accompanying the catalogue were used for the lineage-confirmed gates.

## Results

### The panel passes all reference-genome assertions

The assertion suite passes 24 of 24, including the inverted *S. wolfei* assertion and every negative assertion on *E. coli*.

### No methanogenesis route is assigned to any non-methanogen

The panel was applied to the 1,401 digester MAGs (Table 2).

**Table 2.**
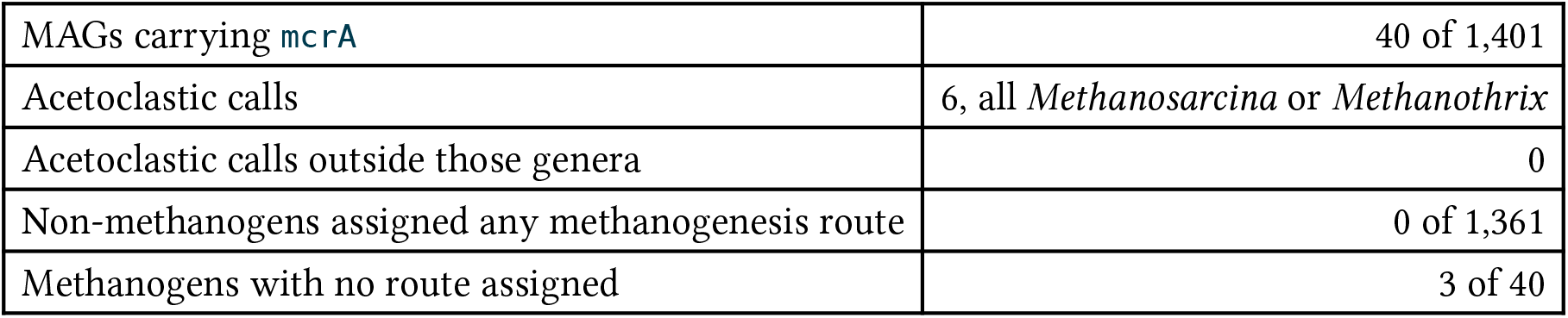
Methanogenesis route assignment across 1,401 digester MAGs.

|  |  |
| --- | --- |
| MAGs carrying *mcrA* | 40 of 1,401 |
| Acetoclastic calls | 6, all *Methanosarcina* or *Methanotherix* |
| Acetoclastic calls outside those genera | 0 |
| Non-methanogens assigned any methanogenesis route | 0 of 1,361 |
| Methanogens with no route assigned | 3 of 40 |

The absence of route assignments among 1,361 bacterial genomes is the principal specificity result. The three methanogens without a route lost it to the lineage policy and the step-based hydrogenotrophic gate: two *Methanothrix* bins carry the C1 pathway but not the acetate markers, and a *Methanothermobacter* bin has three of the six C1 steps. Their earlier hydrogenotrophic calls were the error the policy removes, so they count against sensitivity rather than specificity. Catalogue taxonomy here is the authors’ NCBI names, which give a genus only.

The benchmark also identified a genuine gap. Seven `mcrA`-carrying genomes initially received no route; all carried none of the eight C1 carriers, which is correct biology for the obligate methylotrophs among *Methanofastidiosia* and *Thermoplasmatales*, whose substrate-specific methyltransferases have no licence-clean model. Adding the terminal methylcobalamin:CoM methyltransferase, which detects methylotrophy without naming the substrate, resolved all seven. That model is subfamily-level and matches 30 bacterial genomes on its own; it is sound only because the gate independently requires `mcrA`. Among the digester methanogens, it discriminated correctly, but the GTDB sample showed that it alone also matches non-methylotrophic methanogens, which is why a call resting on it needs a confirming lineage (Methods).

### The export test reduces attributed hydrolytic capacity

Applying the per-protein export test across the same 1,401 genomes (Table 3, Figure 1):

**Table 3.** Genomes carrying a hydrolase family versus those encoding it on the same polypeptide as an export module.

| module | carry the family | export it |  |
| --- | --- | --- | --- |
| Protein hydrolysis | 868 | 106 | 12.2% |
| Carbohydrate hydrolysis | 597 | 50 | 8.4% |
| Lipid hydrolysis | 129 | 7 | 5.4% |
| All intracellular modules | up to 1,337 | 0 | 0.0% |

**Figure 1.**
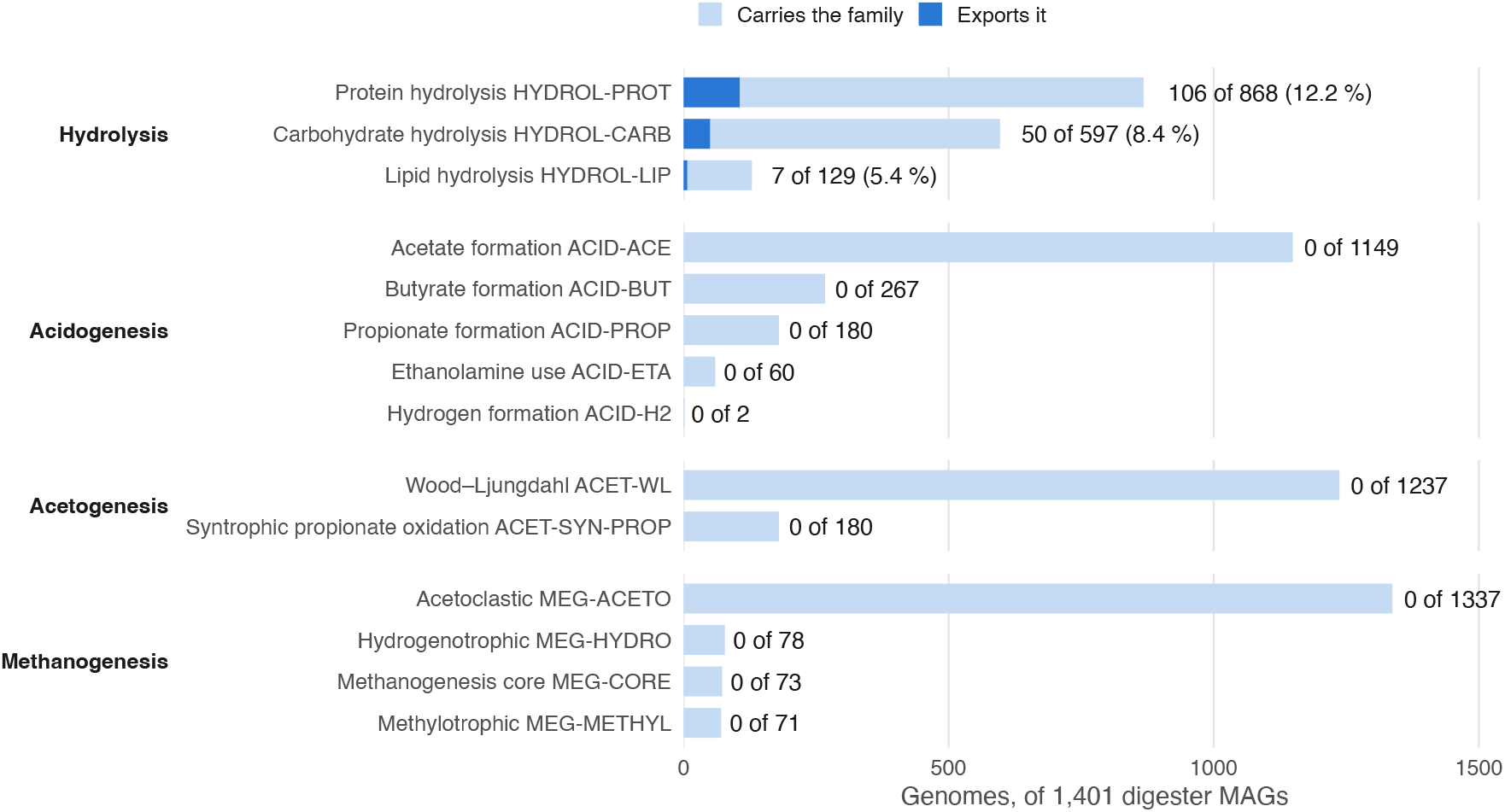
Genomes among the 1,401 digester MAGs that carry at least one core marker of each module (light) and those in which a core marker’s catalytic domain sits on the same polypeptide as an export module (dark). Only the hydrolysis modules have an exported marker. Intracellular modules are counted by any core marker, and some of these are common in bacteria, such as acetate kinase and phosphotransacetylase in the acetoclastic module; carrying one is not a route call.

The zero across every intracellular module is the control: the test fires only on hydrolysis, the sole exported step.

### Specificity holds on genomes built a different way

Applying the panel to the 3,043 GTDB representatives tests whether the digester result depends on one binning pipeline (Table 4). Genomes are classed by the sampling strata with two sourced corrections: *Methanonatronarchaeales* are methyl-reducing methanogens (Sorokin et al., 2017) that the sampling list had counted as background, and genomes in anaerobic methane and alkane oxidiser lineages form a class of their own. One *Methanonatronarchaeales* genome therefore moves from background to methanogen, and 90 oxidisers (89 sampled under methanogen orders, one as background) move to their own class, which is why the classes hold 1,056 and 1,897 genomes rather than the sampled 1,144 and 1,899 (see Methods). Two background genomes were assigned a methanogenesis route, both mcrA-carrying archaea (*Hadarchaeales* and an unnamed order within *Syntropharchaeia*); no bacterial genome received one. On the conservative reading, they are false positives, at a rate of 0.1% in 1,897 background genomes; if the metabolism of these two lineages is established, they belong in the lineage policy with a source, not in the benchmark.

**Table 4.** Methanogenesis route assignment across 3,043 GTDB species representatives.

|  | background | methanogen | anaerobic methane /<br>alkane oxidiser |
| --- | --- | --- | --- |
| genomes | 1,897 | 1,056 | 90 |
| route assigned | 2, both *mcrA*-positive<br>archaea | 819 | 0 |
| no route | 1,895 | 237, of which 217 lack<br>*mcrA* | 90 |

The 90 anaerobic methane and alkane oxidisers (*Methanoperedenaceae, Methanocomedenaceae, Methanogasteraceae*, ANME-1, “*Ca*. Ethanoperedens”) carry mcr and run the pathway in reverse. They are the clearest case for lineage confirmation: 65 of the 90 satisfy the hydrogenotrophic gene gate and 55 the acetoclastic one, because reverse methanogenesis uses the same enzymes. Gene content cannot separate them from methanogens. That none of them receives a route is not an independent test: the class is defined by lineage, and the same lineage policy removes the calls. The one outcome not fixed by construction is the methylotrophic gate, which no oxidiser triggered. Of the 1,056 methanogens, 819 were assigned at least one route. Of the 237 that were not, 217 carry no detectable mcrA at all. GTDB representatives include many incomplete genomes, and a genome that does not contain the defining marker should not be given a route. Sensitivity is therefore 78% of the methanogens sampled, and 98% (819 of 839) of those in which mcrA was detected; most of the gap is genomes that lack the defining marker. The 20 that carry mcrA without an assignable route are the informative residue. They include three *Methanomassiliicoccales*, obligate hydrogen-dependent methylotrophs that lack the Mtr/H4MPT set as a matter of biology and here carry none of the three methyltransferases the methylotrophic gate accepts, neither the substrate-specific MtaB and MttB nor the shared MT2; five *Methanothrix* genomes that carry the C1 pathway but not the acetate markers; and hydrogenotrophs that reach only part of the C1 pathway. Whether a missing marker reflects biology or an incomplete assembly is not resolvable from gene content; in a report, genome completeness (CheckM2) is shown beside every methanogen for that reason.

### A methanogen gets the right route only with a confirming lineage

Specificity against non-methanogens does not show that a methanogen was given the right route. We therefore tabulated every route call by GTDB lineage (`scripts/ summarise_gtdb_benchmark.py`), first with a family-level acetoclastic check and gene-only hydrogenotrophic and methylotrophic gates, then with the lineage policy described in Methods (Table 5, Figure 2).

**Table 5.**
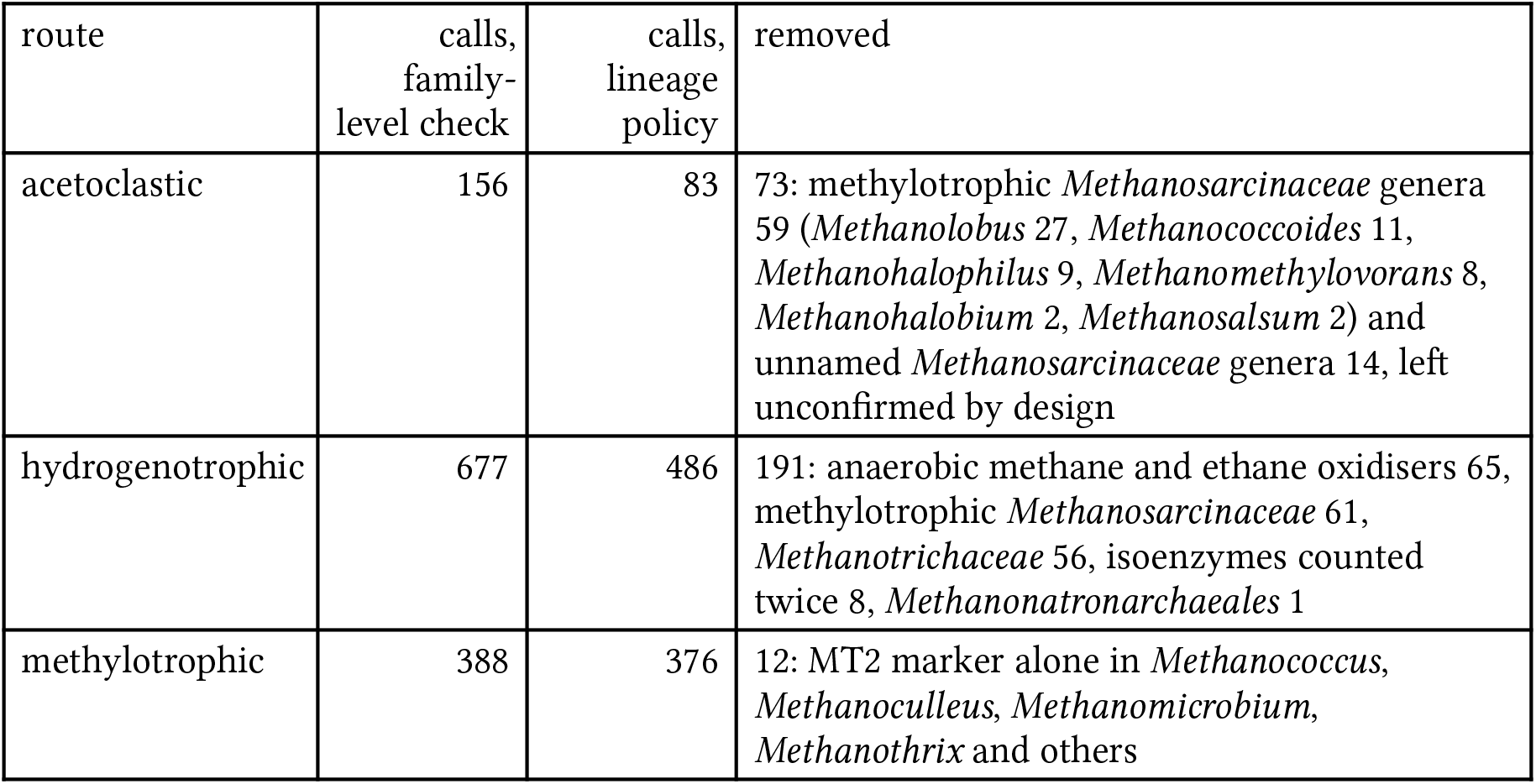
Route calls in the GTDB sample before and after lineage confirmation. After it, every acetoclastic call is *Methanosarcina* (32), *Methanothrix* (29), *Methanocrinis* (12) or an unnamed *Methanotrichaceae* genus (10).

**Figure 2.**
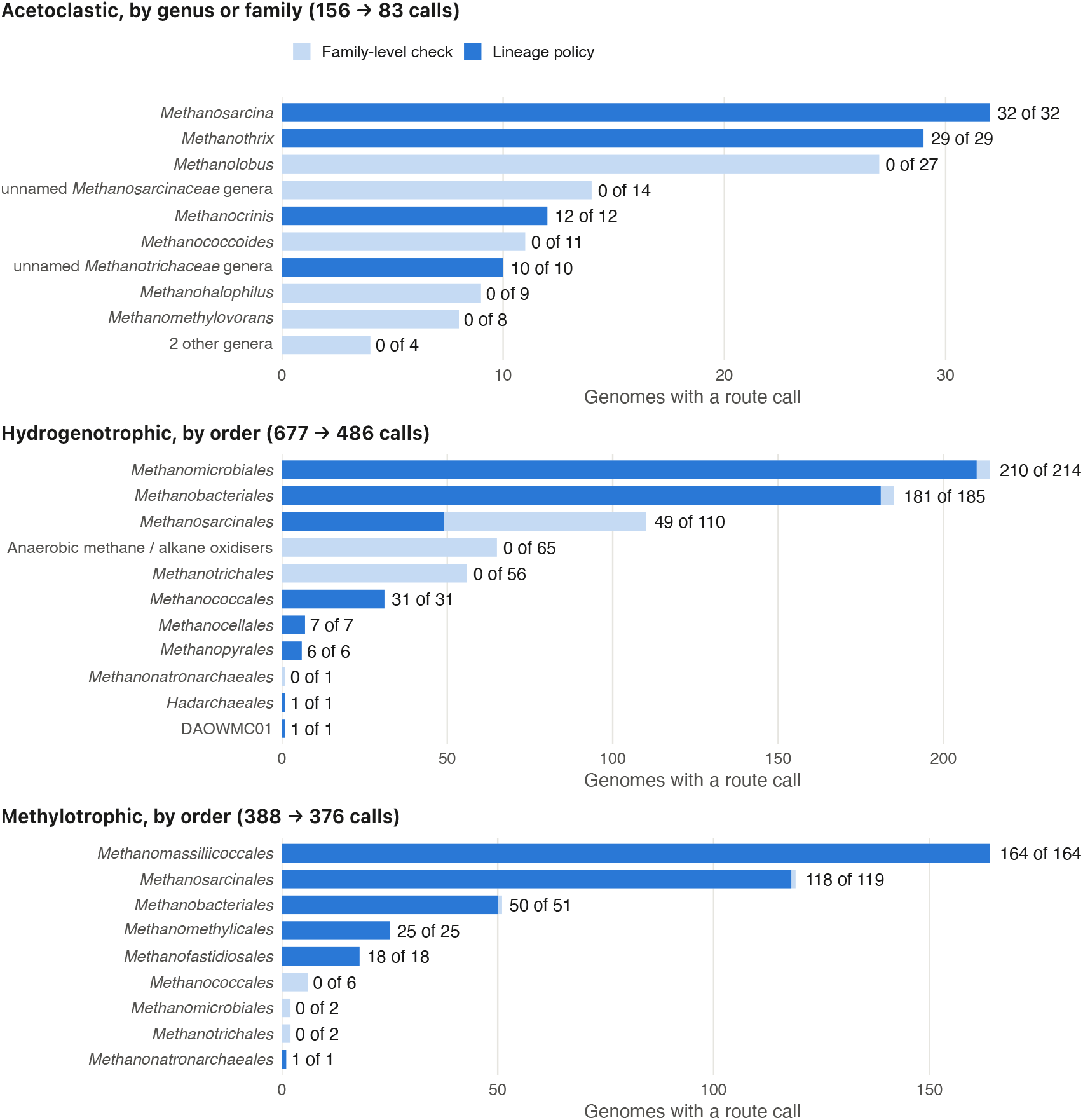
Methanogenesis route calls in the 3,043-genome GTDB sample by lineage, under the family-level acetoclastic check with gene-only hydrogenotrophic and methylotrophic gates (light), and under the lineage policy (dark). Labels give calls kept of calls made. The acetoclastic route is confirmed at genus level (at family level for *Methanotrichaceae*) and the other two at order level, so lineages are shown at genus and order. Anaerobic methane and alkane oxidisers are shown as one group rather than inside the orders that contain them. GTDB placeholder genera are pooled by family, and named acetoclastic genera with at most two calls, all removed, are pooled. The two single-genome hydrogenotrophic rows that are kept, *Hadarchaeales* and DAOWMC01, are the two background genomes counted as false positives in Table 4.

Of the 156 acetoclastic calls under the family-level check, 59 were in methylotrophic genera that do not use acetate and 14 in unnamed *Methanosarcinaceae* genera; the policy leaves them unconfirmed, and more than a quarter of the hydrogenotrophic calls fell on organisms that run the C1 pathway oxidatively or in reverse, or reduce methyl groups rather than CO2 (Sorokin et al., 2017). Neither error is visible in a specificity test against non-methanogens. In the digester catalogue this error did not arise, but only by chance: its six acetoclastic calls all fell in *Methanosarcina* and *Methanothrix*, and the family-level check would have accepted a *Methanolobus* genome just as readily. Most of the change in the number of genomes with a route is correctness, not lost sensitivity (Table 6): only the eight hydrogenotrophs that reach three of the six C1 steps once isoenzymes are counted once are a cost.

**Table 6.** Where the change in route assignments comes from.

| genomes in methanogen orders with $\geq 1$ route | genomes | kind |
| --- | --- | --- |
| family-level check, original order labels | 896 |  |
| anaerobic methane / alkane oxidisers, route removed | -65 | correctness |
| *Methanothrix* genomes whose only call was a false hydrogenotrophic one | -5 | correctness |
| hydrogenotrophs at three of six C1 steps | -8 | sensitivity |
| *Methanonatronarchaeales* moved from background | +1 | label |
| lineage policy, corrected order labels | 819 |  |

Of 50 *Methanobacteriales* genomes with a methylotrophic call, 14 are *Methanosphaera*, confirmed by lineage, and 36 are in genera where methylotrophy is not established (*Methanobacterium* 21, *Methanocatella* 7, *Methanobinarius* 4, *Methanovirga* 2, *Methanarmilla* 2). These calls do not rest on a lone methyltransferase-like hit: all 36 carry the methanol methyltransferase (MtaB) together with a corrinoid protein; 35 of them also carry the methyltransferase partner (MtaA). The corrinoid protein is of the MtaC2 family rather than the MtaC of *Methanosarcina*. The gate requires the corrinoid partner for the MtaB term, and all 36 calls meet that requirement. We report them as capacity and flag them.

### Contig-level co-occurrence does not transfer to the methanogenesis gates

Requiring every gate term on one contig removed every route from 64% of the methanogens that had one under genome-level scoring (525 of 819). The loss grew with the number of contigs in the genome (Figure 3): none for single-contig genomes, 14% at two to ten contigs, 62% at eleven to fifty, 83% at fifty-one to two hundred, and 95% above two hundred. The cause is gene arrangement, not assembly quality: the rule is cheap for an operon-encoded pathway, whose genes sit side by side, and prohibitive for one spread across the chromosome. The mcr, fwd and mtr loci lie apart on the chromosome, so a contig that holds mcrA and four C1 steps has to span most of a replicon.

**Figure 3.**
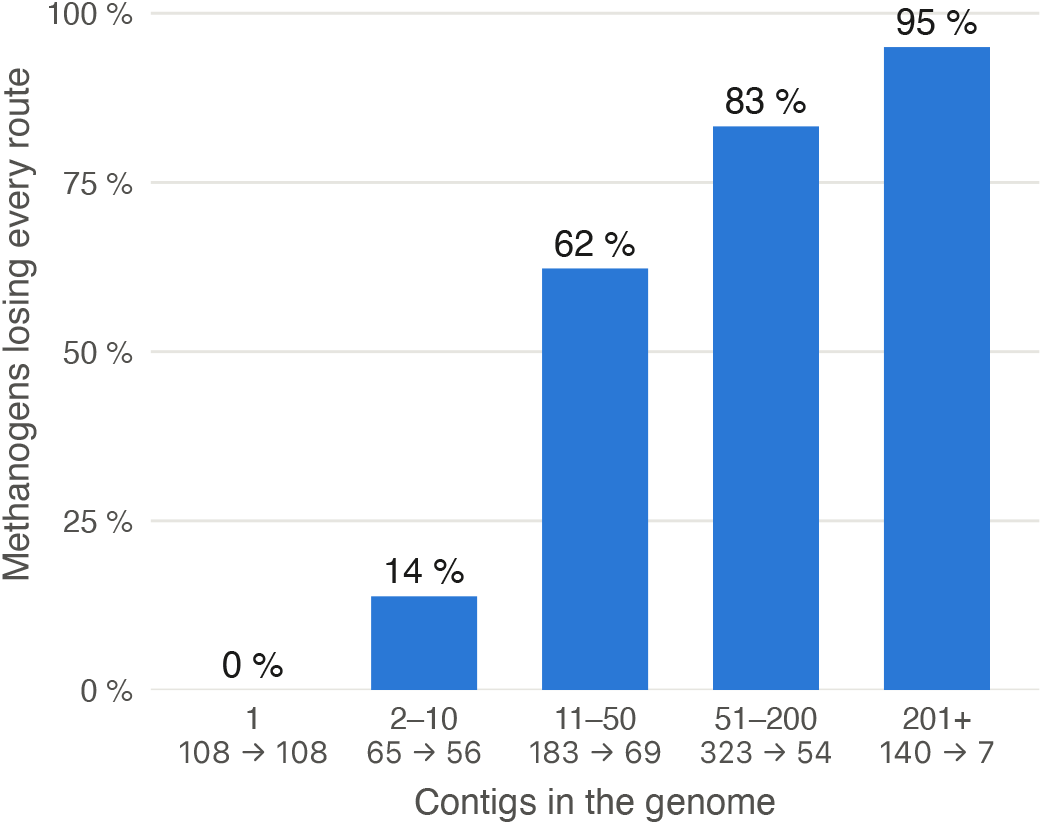
Methanogens of the GTDB sample that lose every route when every gate term must lie on one contig, by the number of contigs in the genome. The share is of genomes with at least one route under genome-level scoring, not of individual route calls. Below each bin, methanogens with a route under genome-level → contig-level scoring. Both scorings use the final scorer and lineage policy; contig counts are taken from the Prodigal gene calls.

### A profile reports the community together with three measurement choices

The numbers in this subsection are properties of the graph as stated, whose arrows are assumptions drawn from the operating handbooks and from how the method works; they are not measurements. The graph (Figure S1) has 28 nodes and 44 arrows; Figure 4 shows the part that concerns the profile. The profile has four direct causes: the community and the three measurement choices (sampling point and time, DNA extraction and sequencing depth, and reference genomes). The reactor state reaches the profile only through the community. Thirteen nodes are causes of the community, directly or through other nodes. Every lever reaches the profile only through the community; the reactor design also reaches it through the sampling point.

**Figure 4.**
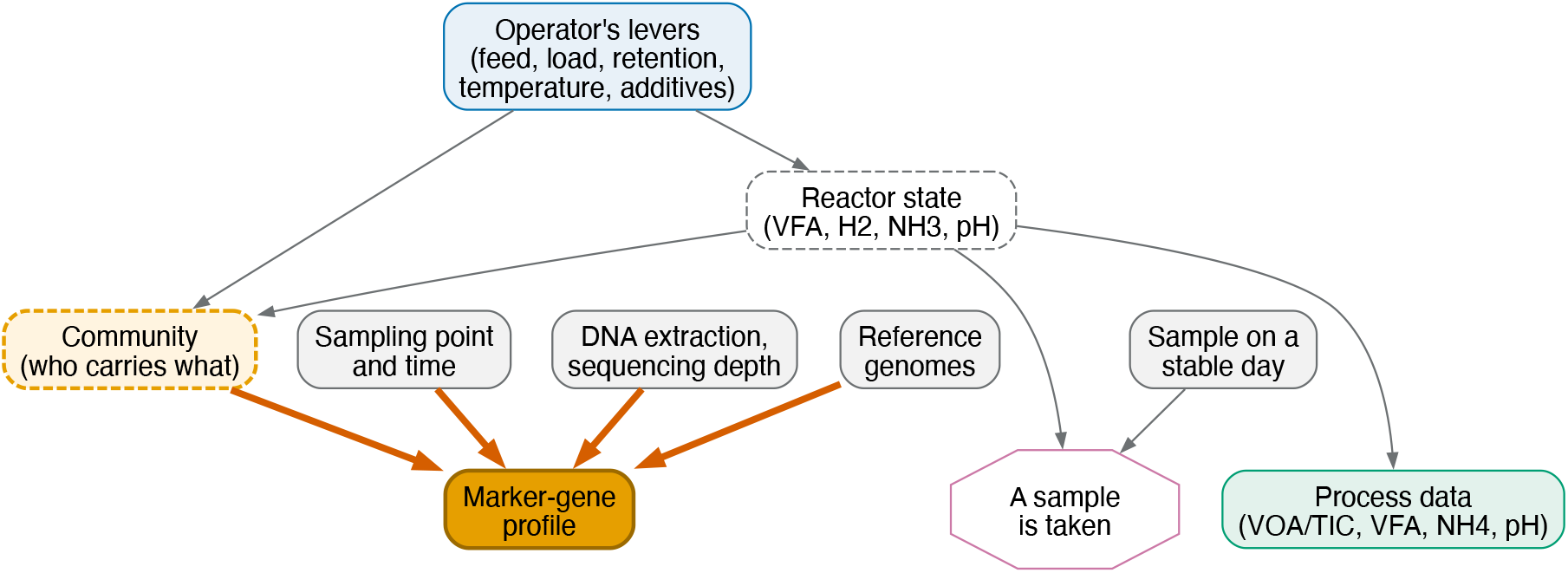
The part of the causal graph that concerns the profile (the full graph is Figure S1). The four thick orange arrows are the direct causes of the profile: the community, the sampling point and time, the DNA extraction and sequencing depth, and the reference genomes. The levers and the reactor state reach the profile only through the community; the process data observe the state directly. Dashed nodes are not observed directly. “A sample is taken” marks selection: analysis is possible only for samples someone chose to take, so every analysis is conditional on it, and samples taken on stable days counter the bias toward disturbed states. Every arrow here is a directed path in the full graph.

The graph implies 24 conditional independencies that involve a measurement. Twelve still hold among samples that were taken, so they can be tested on collected samples. Selection opens the other twelve: they will fail on collected samples even if the graph is right. In the graph, whether a sample is taken depends on the reactor state, on the previous profile, and on whether samples are also taken on stable days. Of the twelve testable ones, nine concern the measurement choices; for example, extraction is independent of the process data, with or without selection. The other three concern the inoculum and the reactor design, and each would fail if one specific arrow that the graph leaves out were present.

For the effect of the operator’s next change on the reactor state in the following interval, the graph has 133 back-door paths, 79 of them open, meaning that they carry association unless something on them is adjusted for. As drawn, no adjustment set exists, because what the operator sees on site drives the decision and is not recorded. If the reason for each change is recorded, the direct causes of the decision (the profile, the process data, the gas yield and the recorded reason) form a sufficient adjustment set.

### Detecting a pyrrolysine marker depends on the gene caller

Monomethylamine methyltransferase illustrates a failure mode that is upstream of the panel entirely. In *Methanosarcina barkeri*, the NCBI reference annotation contains no monomethylamine methyltransferase, so the Pfam family PF05369 returns nothing. Calling genes with Prodigal recovers it. Under the standard translation table, each of the two gene copies is truncated at the in-frame amber codon into fragments of 238 and 202 residues, against a 450-residue model, and both clear the gathering threshold independently. Under a table that reads TAG through, each copy is reconstructed as a single protein of 556 and 535 residues at roughly twice the bit score, with the substituted residue at the expected pyrrolysine position. Across the GTDB sample, the family was found in 278 of 1,144 methanogens, concentrated in *Methanomassiliicoccales* (137), *Methanosarcinales* (119) and *Methanomethylicales* (22), against 18 of 1,899 background genomes. Those background hits cannot by themselves place a genome on a methylotrophic route, because that gate independently requires mcrA.

### Scoring takes seconds per genome

Scoring the 1,401-genome catalogue on 16 cores took 8 min 8 s of wall-clock time and 1.6 CPU-hours in four repeat runs (range 7 min 38 s to 8 min 31 s), or approximately 4 CPU-seconds per genome, of which `hmmsearch` against the 144-model database is nearly all. Peak memory stayed within 16 GB for the job as a whole. The cost scales linearly with genome count and the work is embarrassingly parallel, so a single core processes a genome in a few seconds and a laptop handles a few hundred genomes in minutes; nothing here requires a cluster.

## Discussion

Every methanogenesis gate carries a mandatory `mcrA` term, and `mcrA` is detected only by a gene-specific model assigned by curated accession, so a shared gene symbol or EC elsewhere in the panel cannot by itself place a bacterium on a methanogenesis route. The gates are structurally protected in a way that module completeness is not: a collision inflates a percentage quietly, whereas firing a gate requires the one marker that is hardest to match by accident. This is a design choice, and it is the reason the audits described in Methods, of curated accessions and of symbol matches, are aimed at completeness figures rather than at the gates.

Scoring 1,361 bacterial genomes and assigning none of them a route tests that design directly, and the GTDB representatives extend it: those genomes were not binned at all, so agreement between the digester and GTDB collections cannot be explained by a shared assembly pipeline. The genomes outside the methanogen orders that did receive a route are the informative cases rather than the failures. Each carries `mcrA`, and one belongs to an order of methyl-reducing methanogens that our own list of methanogen orders had omitted, so the panel identified a methanogen before the reference labels did; the benchmark classes were corrected to count it as one. The mcrA requirement protects route calls only. Module completeness percentages have no such required marker, so a gene-symbol or EC collision can raise them unnoticed; the audits in Methods exist for that reason.

The secretion result carries the larger practical consequence. Reporting family presence alone would credit 597 genomes with cellulolytic capacity where 50 is defensible, in the module most often identified as rate-limiting for fibrous feedstocks. An operator told that 43% of a community is cellulolytic will draw different conclusions than one told the figure is 4%, and only the second is supported by the evidence. The zero across intracellular modules indicates the test is measuring export architecture rather than domain co-occurrence generally.

The finding that seven mcrA-carrying genomes initially received no route illustrates the value of reporting non-measurement explicitly. Had those genomes been scored as zero rather than as unassigned, the gap would have read as a biological result about *Methanofastidiosia* rather than as a missing detector, and would probably not have been investigated.

### From profile to advice

The causal graph (Figure 4) turns two caveats into consequences of its structure. First, the reactor state reaches the profile only through the community, which changes over weeks. A profile therefore cannot report what the reactor is doing now; the capacity limitation below follows from the graph. Second, three measurement choices act on the profile directly, so a change between two samples can be read as a change in the community only if sampling, extraction and reference stay the same. Because extraction stays independent of the process data, a practical rule follows: if the profile moves while volatile fatty acids, VOA/TIC and gas yield do not, check the protocol before the community.

A single finding cannot be read on its own either. Thirteen nodes act on the community, so a thin acetoclastic route is compatible with ammonia inhibition, trace-element limitation, washout, or a route that was never established. The reactor state also acts on the community, so a thin acetate route can be a result of high ammonia, which inhibits acetate-using methanogens, rather than the cause of a problem such as acetate accumulation. Interpreting a profile therefore needs a second measurement of a competing cause, such as the process data or a digester trace-element analysis. The profile complements the physicochemical proxies; it does not replace them.

Whether advice based on profiles improves outcomes is a causal question, and the graph shows what a prospective validation would have to record to answer it (Maiväli, 2026). The estimand, the precise effect to be estimated, has to be one intervention. An example is raising the loading rate by one step rather than holding it. The outcome would be a process upset within eight weeks, read from the plant’s routine log as VOA/TIC above that plant’s own upper limit. Advice based on a profile claims something more specific: that the same load step is riskier in a reactor whose acetate route is thin than in one whose route is strong. In causal terms, the community modifies the effect of the load step (effect modification). Testing that advice therefore needs more than showing that load steps cause upsets: the effect has to be larger in reactors the profile flags as thin than in those it does not.

Operators also act on what they see on site, so the effect is confounded by indication, and without a record of why each change was made no adjustment set exists. A prospective validation therefore has to record the reason for each change at the time, on a fixed checklist, together with the profile, the process data and the gas yield. Framing the comparison as an emulated target trial (Hernán & Robins, 2016), with time zero at the delivery of the profile, places each plant in a group (raise the load or hold it) by what was decided at that point. Grouping plants by what they did in the following weeks would put every plant that had an upset before it could raise the load into the “hold” group, and so make raising the load look safer than it is. The comparison is impossible wherever all operators respond to a signal in the same way, because there is then no one to compare with (positivity fails).

Selection, analysing only the samples someone chose to take, matters as much as confounding: it biases the sample rather than merely thinning it. Plants sample when a process is in difficulty; on such samples, half of the graph’s implied independencies cannot be tested, and thresholds calibrated on them would describe disturbed states. Samples taken on stable days are the counter-weight. This is what the prospective validation named under benchmark scope below would have to look like.

### Limitations

Capacity, not activity. The panel measures what a community is equipped to do. It cannot report expression, rate or flux; those require transcriptomics, proteomics or process measurement. Secretion evidence extends the claim from *encoded* to *encoded and built for export*, which is still not *active*.

Markers with no licence-clean detector. Nine of 86 markers remain unresolved. bcd and crt have no model in either source, so syntrophic butyrate oxidation is reported as unassessable rather than absent, including in an organism that performs it.

Dependence on upstream gene calling. The pyrrolysine result above is a property of the annotation, not of the panel: `mtmB` is detectable from Prodigal calls and invisible in NCBI reference proteomes. Any input built by an annotation pipeline that does not translate through in-frame amber codons will under-report methylamine methyltransferases, and the panel cannot detect that this has happened. The same caution applies to any marker whose gene carries a recoded codon. Family-level resolution in hydrolysis. Pfam families do not distinguish secreted from cytoplasmic enzymes on their own (hence the export test), nor do they resolve substrate specificity within a family. The export layer detects the cellulosomal and cell-surface-anchored routes; a plain Sec-secreted enzyme carrying no binding module would be missed.

Taxonomy dependence. The acetoclastic, hydrogenotrophic and MT2-only methylotrophic calls are only as good as the supplied lineage and the policy that reads it. Without taxonomy they degrade to gene evidence, which over-calls, and the NCBI names that accompany many catalogues give a genus only, too coarse for the order-level rules. The policy is data, one table with a source per row, so a new lineage is a reviewed edit rather than a code change; a lineage it does not know is reported, not guessed.

Prevalence is measured over binned genomes. Scoring is per genome, so a marker carried by taxa that do not bin is not counted. The error always lowers prevalence, and it falls hardest on rare or taxonomically restricted functions, whose carriers are the least likely to form bins.

Scoring assembly proteomes directly would recover the unbinned carriers, but it is not a safe substitute here, for a quantitative rather than a conceptual reason. A single-gene call survives the move, because presence in a sample does show that some organism carries the gene. A multigene gate does not: different organisms can satisfy its members, and the false-positive rate grows with how common the individual markers are. The hydrogenotrophic gate is the worst case in this panel. Its threshold of four present among eight carriers lets a call be assembled from unrelated genomes in many ways.

Contig-level co-occurrence keeps the evidence within one organism without a bin, but it does not transfer to these gates (Figure 3). The option remains for pathways it suits; these gates are defined for binned genomes and should be run on them. Binned input answers what an organism can do, and assembly-level search answers whether a sample contains a capability. The two questions have different error models and should be kept apart.

Taxonomic ground truth. Both the specificity and the sensitivity figures are measured against order-level labels, and those labels were wrong in both directions here. *Methanonatronarchaeales* are methanogens that our order list did not include, so a correct call counted as a false positive until the benchmark classes were corrected. The anaerobic methane and alkane oxidisers sit inside methanogen orders; an order-level benchmark counts their routes as correct methanogen calls, and it did so for 65 of them before the lineage policy. Either way, an error in the reference labels appeared as an error of the panel, so a specificity test against non-methanogens has to be paired with a check of which route each methanogen receives.

Benchmark scope. Specificity rests on collections built differently, one binned from digester metagenomes by a single group and one drawn from GTDB representatives, which removes the possibility that the result is an artefact of one binning pipeline. Both remain snapshots of public sequence databases, and neither tests genomes from a digester the panel has not seen. No prospective validation in operating plants has been attempted; what one would have to record is set out in the Discussion. Sensitivity is measured only against taxonomic assignment, which is itself an inference: a genome placed in a methanogen order that carries no mcrA is counted here as a genome with a missing marker, not as a misplaced lineage, and the two are not distinguishable from gene content alone.

### Outlook

The markers that remain unresolved are not evenly distributed: they concentrate in syntrophic fatty-acid oxidation and in substrate-specific methylotrophy, both of which matter for interpreting a stressed reactor. Closing them requires new licence-clean models. The pyrrolysine case suggests where else to look: the obstacle there was never the model but the annotation feeding it, and any marker whose gene carries a recoded codon will behave the same way. The approach itself is not specific to anaerobic digestion; the same construction, a curated domain-scoped pathway model over public-domain and CC0 profile HMMs, applies to any setting where licensing rather than biology is the obstacle.

### Data and code availability

Panel, scorer, build and validation scripts are distributed as the `digestome` package. They run as standalone command-line steps; the scorer and aggregator are Python 3 standard library only, so they carry no dependency beyond HMMER and a Python interpreter. Reference data are downloaded by the provided scripts from NCBIfam, InterPro/Pfam and Rhea; no licensed database is required at any stage. The benchmark catalogue is available from NCBI BioProject PRJNA602310 (Campanaro et al., 2020). Code is released under the MIT licence and the curated panel under CC BY 4.0.

Every number in this manuscript was produced with version 0.1.2 (Git tag v0.1.2). Releases are archived at Zenodo under DOI 10.5281/zenodo.22913615, which lists every version and resolves to the latest, and the code is developed at https://github.com/tpall/digestome.

## Competing interests

The author is the owner of Magrittr OÜ, which commercially provides anaerobic digestion microbiome analyses using the panel described herein. The panel, scoring methodology and validation scripts are openly available to facilitate independent replication and verification of the results.

## Funding

No external funding. Computation was bought from the University of Tartu High Performance Computing Centre under a commercial service agreement.

## Acknowledgements

The author thanks the High Performance Computing Centre of the University of Tartu for its computing service (University of Tartu, 2018), and the authors of the biogas MAG catalogue (Campanaro et al., 2020) for making both the genomes and the per-sample abundances public: the specificity benchmark in this work is only possible because they did.

## Supplementary figure

**Figure S1.**
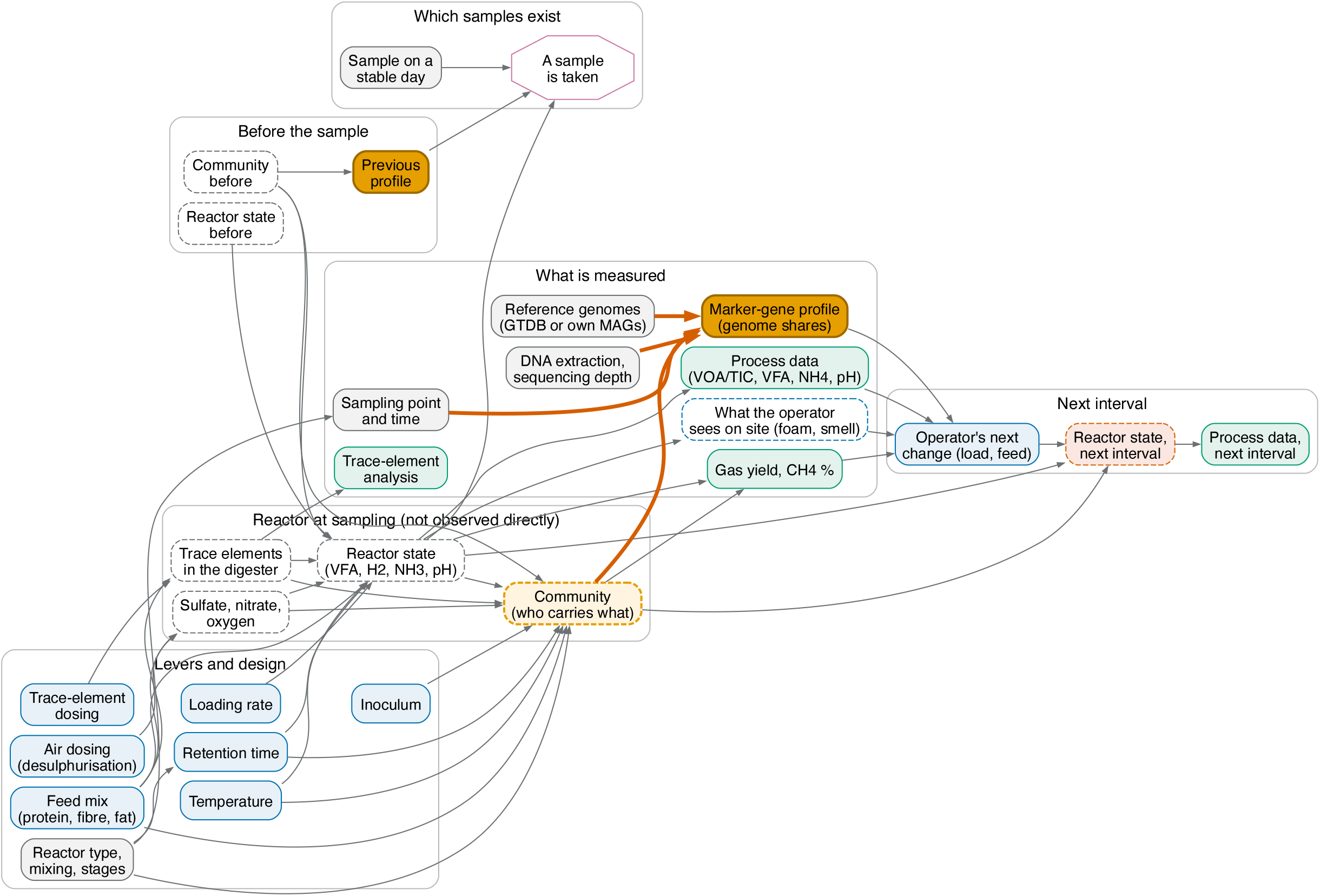
Full causal graph of a digester and of what a marker-gene profile observes, unrolled into three time slices: the weeks before the sample, the sampling week, and the eight weeks after the profile is delivered. Dashed nodes are not observed directly. The four thick orange arrows are the direct causes of the profile. “A sample is taken” marks selection: analysis is possible only for samples someone chose to take, so every analysis is conditional on it. The script that draws the figure gives a reason for every arrow (`manuscript/dag/`). The figure is turned 90 degrees.

## Notes

https://doi.org/10.5281/zenodo.22913615

https://github.com/tpall/digestome

https://www.ncbi.nlm.nih.gov/bioproject/PRJNA602310

https://data.gtdb.ecogenomic.org/releases/release226/

### Competing Interest Statement

The author is the owner of Magrittr OÜ, which commercially provides anaerobic digestion microbiome analyses using the panel described herein.

### Summary of Updates

Version 2 adds an analysis of what a marker-gene profile can and cannot attribute. A causal graph of a digester and of its measurement (new Methods subsection, new Results subsection, Figure 4, and the full graph as Figure S1) shows that a profile reports the community together with three measurement choices (sampling, DNA extraction and sequencing depth, and reference genomes). The Discussion interprets this and states what a prospective validation in operating plants would have to record. The contig-level co-occurrence analysis moved from the Limitations into the Results (Figure 3). The abstract has one new sentence, and "profile" is defined at its first use. Figures from a separate, uncited bile-acid analysis were removed from the Limitations; the argument they illustrated stays. No benchmark number of version 1 changed.

## References

AFMB-CNRS, & University of Nebraska-Lincoln. (n.d.). CAZy and dbCAN terms of use. https://www.cazy.org/, https://bcb.unl.edu/dbCAN2/. Retrieved August 10, 2026, from https://www.cazy.org/

Campanaro, S., Treu, L., Rodriguez-R, L. M., Kovalovszki, A., Ziels, R. M., Maus, I., Zhu, X., Kougias, P. G., Basile, A., Luo, G., Schlüter, A., Konstantinidis, K. T., & Angelidaki, I. (2020). New insights from the biogas microbiome by comprehensive genome-resolved metagenomics of nearly 1600 species originating from multiple anaerobic digesters. Biotechnology for Biofuels, 13(1), 25. 10.1186/s13068-020-01679-y

Carballa, M., Regueiro, L., & Lema, J. M. (2015). Microbial management of anaerobic digestion: Exploiting the microbiome-functionality nexus. Current Opinion in Biotechnology, 33, 103–111. 10.1016/j.copbio.2015.01.008

Chaumeil, P.-A., Mussig, A. J., Hugenholtz, P., & Parks, D. H. (2022). GTDB-Tk v2: Memory friendly classification with the Genome Taxonomy Database. Bioinformatics, 38(23), 5315–5316. 10.1093/bioinformatics/btac672

De Vrieze, J. (2020). The next frontier of the anaerobic digestion microbiome: From ecology to process control. Environmental Science and Ecotechnology, 3, 100032. 10.1016/j.ese.2020.100032

Eddy, S. R. (2011). Accelerated profile HMM searches. PLoS Computational Biology, 7(10), e1002195. 10.1371/journal.pcbi.1002195

Fricke, W. F., Seedorf, H., Henne, A., Krüer, M., Liesegang, H., Hedderich, R., Gottschalk, G., & Thauer, R. K. (2006). The genome sequence of Methanosphaera stadtmanae reveals why this human intestinal archaeon is restricted to methanol and H2 for methane formation and ATP synthesis. Journal of Bacteriology, 188(2), 642–658. 10.1128/JB.188.2.642-658.2006

Geller-McGrath, D., Konwar, K. M., Edgcomb, V. P., Pachiadaki, M., Roddy, J. W., Wheeler, T. J., & McDermott, J. E. (2024). Predicting metabolic modules in incomplete bacterial genomes with MetaPathPredict. eLife, 13, e85749. 10.7554/eLife.85749

Hahn, C. J., Laso-Pérez, R., Vulcano, F., Vaziourakis, K.-M., Stokke, R., Steen, I. H., Teske, A., Boetius, A., Liebeke, M., Amann, R., Knittel, K., & Wegener, G. (2020). “Candidatus Ethanoperedens,” a thermophilic genus of archaea mediating the anaerobic oxidation of ethane. mBio, 11(2), e00600–20. 10.1128/mBio.00600-20

Haroon, M. F., Hu, S., Shi, Y., Imelfort, M., Keller, J., Hugenholtz, P., Yuan, Z., & Tyson, G. W. (2013). Anaerobic oxidation of methane coupled to nitrate reduction in a novel archaeal lineage. Nature, 500(7464), 567–570. 10.1038/nature12375

Hassa, J., Klang, J., Benndorf, D., Pohl, M., Hülsemann, B., Mächtig, T., Effenberger, M., Pühler, A., Schlüter, A., & Theuerl, S. (2021). Indicative marker microbiome structures deduced from the taxonomic inventory of 67 full-scale anaerobic digesters of 49 agricultural biogas plants. Microorganisms, 9(7), 1457. 10.3390/microorganisms9071457

Hernán, M. A., & Robins, J. M. (2016). Using big data to emulate a target trial when a randomized trial is not available. American Journal of Epidemiology, 183(8), 758–764. 10.1093/aje/kwv254

Jetten, M. S. M., Stams, A. J. M., & Zehnder, A. J. B. (1992). Methanogenesis from acetate: A comparison of the acetate metabolism in Methanothrix soehngenii and Methanosarcina spp. FEMS Microbiology Reviews, 8(3-4), 181–198. 10.1111/j.1574-6968.1992.tb04987.x

Kananen, K., Veseli, I., Quiles Pérez, C. J., Miller, S. E., Eren, A. M., & Bradley, P. H. (2025). Adaptive adjustment of profile HMM significance thresholds improves functional and metabolic insights into microbial genomes. Bioinformatics Advances, 5(1), vbaf039. 10.1093/bioadv/vbaf039

Khomyakova, M. A., Merkel, A. Y., Slobodkin, A. I., & Sorokin, D. Y. (2023). Phenotypic and genomic characterization of the first alkaliphilic aceticlastic methanogens and proposal of a novel genus Methanocrinis gen. Nov. Within the family Methanotrichaceae. Frontiers in Microbiology, 14, 1233691. 10.3389/fmicb.2023.1233691

Knittel, K., & Boetius, A. (2009). Anaerobic oxidation of methane: Progress with an unknown process. Annual Review of Microbiology, 63(1), 311–334. 10.1146/annurev.micro.61.080706.093130

Koch, C., Müller, S., Harms, H., & Harnisch, F. (2014). Microbiomes in bioenergy production: From analysis to management. Current Opinion in Biotechnology, 27, 65–72. 10.1016/j.copbio.2013.11.006

Maiväli, Ü. (2026). The Bayesian way to causal epidemiology. https://ymaivali.github.io/meth_book/

Mistry, J., Chuguransky, S., Williams, L., Qureshi, M., Salazar, G. A., Sonnhammer, E. L. L., Tosatto, S. C. E., Paladin, L., Raj, S., Richardson, L. J., Finn, R. D., & Bateman, A. (2021). Pfam: The protein families database in 2021. Nucleic Acids Research, 49(D1), D412–D419. 10.1093/nar/gkaa913

Nobu, M. K., Narihiro, T., Kuroda, K., Mei, R., & Liu, W.-T. (2016). Chasing the elusive Euryarchaeota class WSA2: Genomes reveal a uniquely fastidious methyl-reducing methanogen. The ISME Journal, 10(10), 2478–2487. 10.1038/ismej.2016.33

Oren, A. (2014). The family Methanosarcinaceae. In The prokaryotes: Other major lineages of bacteria and the archaea (pp. 259–281). Springer. 10.1007/978-3-642-38954-2_408

Ostos, I., Flórez-Pardo, L. M., & Camargo, C. (2024). A metagenomic approach to demystify the anaerobic digestion black box and achieve higher biogas yield: A review. Frontiers in Microbiology, 15, 1437098. 10.3389/fmicb.2024.1437098

Palù, M., Basile, A., Zampieri, G., Treu, L., Rossi, A., Morlino, M. S., & Campanaro, S. (2022). KEMET: A python tool for KEGG Module evaluation and microbial genome annotation expansion. Computational and Structural Biotechnology Journal, 20, 1481–1486. 10.1016/j.csbj.2022.03.015

Pathway Solutions / Kanehisa Laboratories. (n.d.). KEGG licensing. https://www.pathway.jp/en/licensing.html, https://www.kegg.jp/kegg/legal.html. Retrieved September 28, 2026, from https://www.pathway.jp/en/licensing.html

Shaffer, M., Borton, M. A., McGivern, B. B., Zayed, A. A., La Rosa, S. L., Solden, L. M., Liu, P., Narrowe, A. B., Rodríguez-Ramos, J., Bolduc, B., Gazitúa, M. C., Daly, R. A., Smith, G. J., Vik, D. R., Pope, P. B., Sullivan, M. B., Roux, S., & Wrighton, K. C. (2020). DRAM for distilling microbial metabolism to automate the curation of microbiome function. Nucleic Acids Research, 48(16), 8883–8900. 10.1093/nar/gkaa621

Singh, A., Moestedt, J., Berg, A., & Schnürer, A. (2021). Microbiological surveillance of biogas plants: Targeting acetogenic community. Frontiers in Microbiology, 12, 700256. 10.3389/fmicb.2021.700256

Smith, K. S., & Ingram-Smith, C. (2007). Methanosaeta, the forgotten methanogen? Trends in Micro biology, 15(4), 150–155. 10.1016/j.tim.2007.02.002

Sorokin, D. Y., Makarova, K. S., Abbas, B., Ferrer, M., Golyshin, P. N., Galinski, E. A., Ciordia, S., Mena, M. C., Merkel, A. Y., Wolf, Y. I., Loosdrecht, M. C. M. van, & Koonin, E. V. (2017). Discovery of extremely halophilic, methyl-reducing euryarchaea provides insights into the evolutionary origin of methanogenesis. Nature Microbiology, 2(8), 17081. 10.1038/nmicrobiol.2017.81

Textor, J., Zander, B. van der, Gilthorpe, M. S., Liśkiewicz, M., & Ellison, G. T. H. (2016). Robust causal inference using directed acyclic graphs: The R package ‘dagitty’. International Journal of Epidemiology, 45(6), 1887–1894. 10.1093/ije/dyw341

Theuerl, S., Herrmann, C., Heiermann, M., Grundmann, P., Landwehr, N., Kreidenweis, U., & Prochnow, A. (2019). The future agricultural biogas plant in Germany: A vision. Energies, 12(3), 396. 10.3390/en12030396

Theuerl, S., Klang, J., & Prochnow, A. (2019). Process disturbances in agricultural biogas production —causes, mechanisms and effects on the biogas microbiome: A review. Energies, 12(3), 365. 10.3390/en12030365

University of Tartu. (2018). UT Rocket. share.neic.no. 10.23673/PH6N-0144

Vanwonterghem, I., Evans, P. N., Parks, D. H., Jensen, P. D., Woodcroft, B. J., Hugenholtz, P., & Tyson, G. W. (2016). Methylotrophic methanogenesis discovered in the archaeal phylum Verstraetearchaeota. Nature Microbiology, 1(12), 16170. 10.1038/nmicrobiol.2016.170

Zhou, Z., Tran, P. Q., Breister, A. M., Liu, Y., Kieft, K., Cowley, E. S., Karaoz, U., & Anantharaman, K. (2022). METABOLIC: High-throughput profiling of microbial genomes for functional traits, metabolism, biogeochemistry, and community-scale functional networks. Microbiome, 10(1), 33. 10.1186/s40168-021-01213-8

