## Supplementary figures and images for "digestome: a licence-clean marker-gene panel for functional profiling of anaerobic digestion microbiomes"

### FigureS1

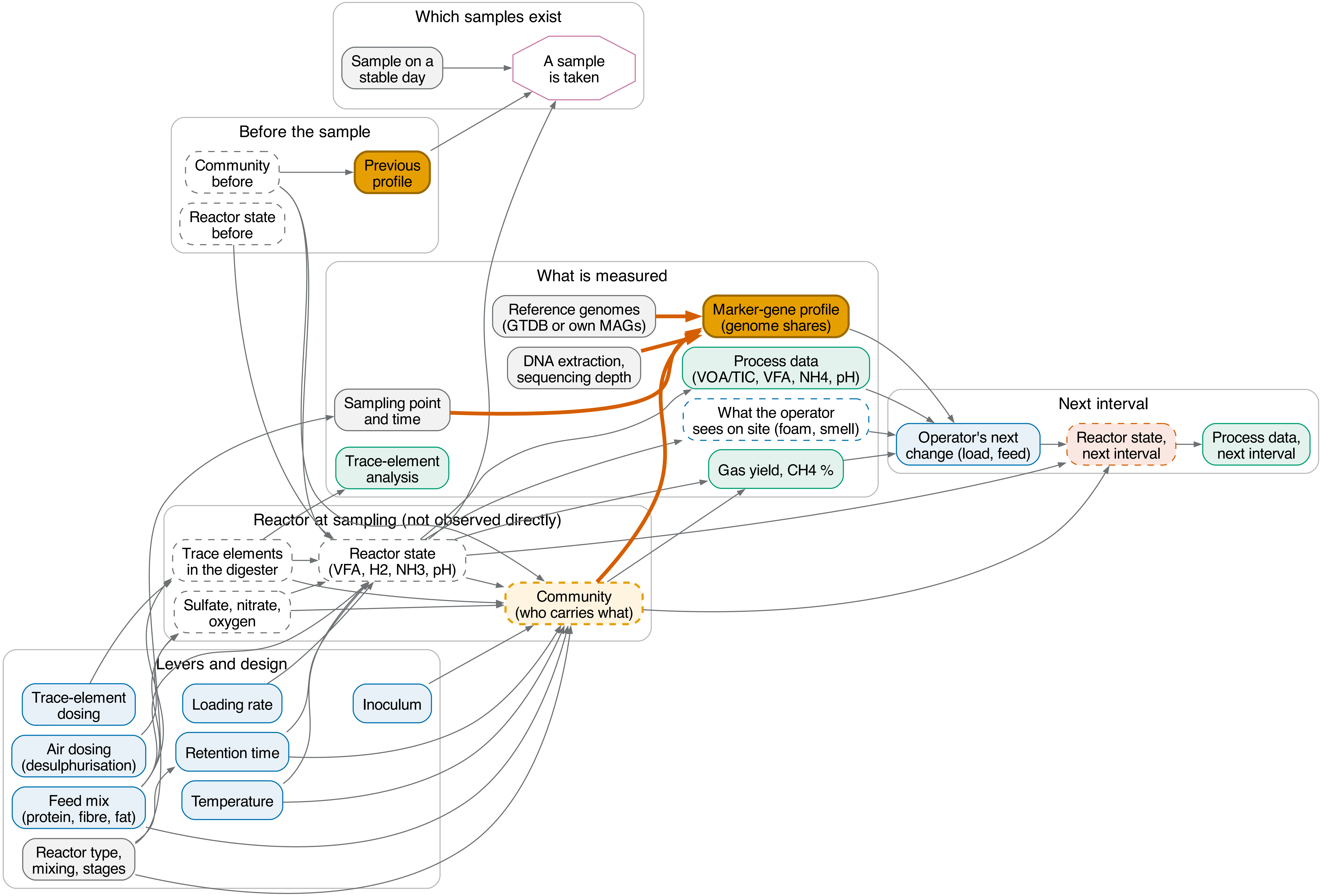
